# The influence of stimulus duration on olfactory identity

**DOI:** 10.1101/2020.09.07.286773

**Authors:** Praveen Kuruppath, Leonardo Belluscio

## Abstract

Duration of a stimulus plays an important role in coding of sensory information. The role of stimulus duration is extensively studied in tactile, visual and auditory system. In the olfactory system, how the stimulus duration influences the identity of an olfactory information is not well understood. To test this, we activated the olfactory bulbs with blue light in mice expressing channelrhodopsin and behaviorally assessed the relevance of stimulus duration on olfactory identity. Our behavior data demonstrate that stimulus duration changes the olfactory information and the associated behavior.

## Introduction

Stimulus duration is an important parameter of the sensory stimuli. The significance of stimulus duration has extensively studied in visual, auditory and tactile sensory system[1–4]. In the olfactory system the role of stimulus duration is not well understood because of difficulty controlling the input duration which is affected by sniffing[5] and because of complex interactions between odorants and the epithelium[6,7]. Previous studies have demonstrated the behavioral relevance of precise olfactory timing relative to sniffing[8,9]. Temporal properties of the stimulus are important for obtaining information about the odor plumes when the odor is delivered into the environment. For mammals, temporal information serves as an important cue for finding the odor source in their natural environment to locate food, mates, and avoid predators[10–15].

For olfaction, temporal information can be significant for identifying an odor. Previous studies in honeybees by Wright and co-workers have observed that increasing the sampling time improves their ability to recognize and differentiate odors[16]. Their results show that sampling time affects olfactory learning, recognition, and discrimination, suggesting that having longer time to sample an odor stimulus yields more information about an odor’s presence and its molecular identity, which improves the ability of the olfactory system to form a neural representation of an odor’s molecular identity. However, in mammals, no studies have explored the behavioral relevance of stimulus duration in the olfactory identity, due to the fact that input duration is widely affected by the sniffing.

In this study we controlled the sensory inputs to the olfactory bulbs (OB) with blue light in tetO-ChIEF-Citrine mice line, where channelrhodopsin-2 is expressed in all the olfactory sensory neurons (OSNs) and studied the behavioral responses to the light pulse stimulation with different stimulus durations. We found that mice respond differently to shorter and longer stimulus durations, suggesting that olfactory information changes with the stimulus duration.

## Materials and methods

### Experimental animals

All animal procedures conformed to National Institutes of Health guidelines and were approved by the National Institute of Neurological Disorders and Stroke Institutional Animal Care and Use Committee. Mice were bred in-house and were maintained on a 12 h light/dark cycle with food and water ad libitum.

The tetO-ChIEF-Citrine line, was generated from pCAGGS-I-oChIEF-mCitrine-I-WPRE (7.7kb; Roger Tsien, UCSD), which contains the coding sequence for mammalian-optimized ChIEF fused to the yellow fluorescent protein Citrine, at the National Institute of Mental Health Transgenic Core Facility (Bethesda, MD) as previously described [17,18]. The OMP-tTA knock-in mouse line expressing the tetracycline transactivator protein (TTA) under the control of the OMP-promoter was a gift from Dr. Joseph Gagos. Experimental animals were OMP-tTA+/- / tetO-ChIEF-Citrine+/- (OMP-ChIEF), generated by crossing heterozygous tetO-ChIEF-Citrine (tetO-ChIEF-Citrine+/-) with homozygous OMP-tTA mice (OMP-tTA-/-).

### Genotyping

OMP-ChIEF pups were identified by the visualization of fluorescence in the nose and OB of P0-P2 pups under epifluorescence illumination.

### Animal preparations

Data was collected from 12 OMP-ChIEF mice. Experimental animals were prepared as described previously[19,20]. Briefly, mice were anesthetized with an intraperitoneal injection of Ketamine/Xylazine mixture 100 and 10 mg/kg body weight, respectively. Each animal was fixed with a stereotactic frame with the head held in place by a bar tie to each temporal side of the skull. The animals were kept warm with hand warmers (Grabber, Grand Rapids, MI, USA). Surgery was started when the animal showed no movement in response to foot pinching. A craniotomy was performed above the skull over each OB. Fiber optic pins were implanted on the dorsal surface of each OB as described previously [19,20]. Mice were injected with Ketoprophen (5mg/kg) immediately after the surgery. Animals were allowed to recover in their home cage for one week.

### Behavioral procedure and training

#### Light stimulation and foot shock avoidance training

Behavioral training began after the animals recovered from the surgery (one week). Training was performed for two days on a modified ‘Y’ maze with two equally sized open arms and one permanently closed arm. Each open arm was independently paved with an electric grid shock floor. The mice were connected to a 400 μm core-diameter optical fiber attached to a 473 nm solid-state variable-power laser (LaserGlow Technologies, Toronto, Canada, Figure. 1A) and allowed to habituate in the ‘Y’ maze for 15 minutes. The time spent in each arm of the ‘Y’ maze was recorded and assessed for arm preference. The preferred arm for the mice was selected as the Light zone and the opposite arm served as the Safe zone. If the mice did not demonstrate a preference for either arm, the Light zone was randomly selected. After the habituation, mice were re-introduced to the ‘Y’ maze for the ten-minute training session. A reinforcement training was performed every day prior to the test session to enhance learning. The training included light stimulation followed by a mild foot shock. The foot shock avoidance training is paired with either left or right OB stimulation. The light stimulation and the foot shock were delivered in the Light zone when the mice completely entered that zone (Figure 1B). The mice had free access to the Safe zone to escape the foot shock. Light stimulation, consisting of a train of ten light pulses of 50 ms duration with an interval of 150 ms, was externally triggered by a Master-8 timer (A.M.P.I, Jerusalem, Israel). The output power of the light pulses was measured and adjusted to 20-22 mW. The mild foot shock (0.65mAmps, 5 s) generated by a stand-alone shock generator (Med Associates, USA) was delivered 2 s after the light stimulation by Master-8 timer. The mice were trained to move to the Safe zone when the light stimuli and the foot shock were delivered in the Light zone.

**Figure 1.**
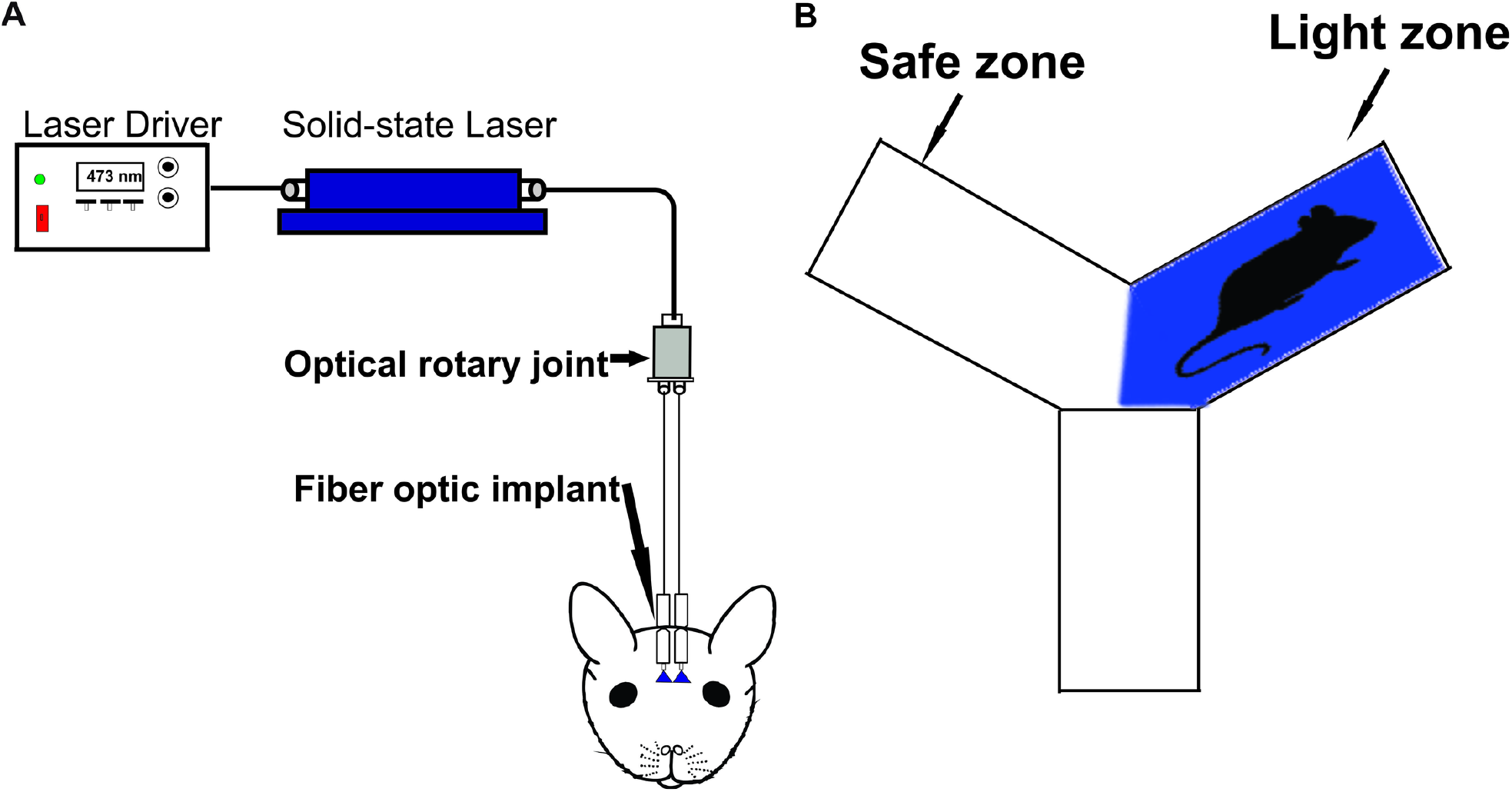
Optogenetic stimulation of olfactory bulb. A, Schematic diagram of the functional system. B, Behavioral setup. Light zone indicates the area where the light stimulation is delivered. Safe zone indicates the area the mice can escape from the light stimulation and foot shock.

### Light zone avoidance test

The Light zone avoidance test was performed for two days. A reinforcement training was performed each day before the testing. Test stimuli, consisting of a train of ten light pulses of 10 or 25 ms duration with an interval of 150 ms. The testing began 60 minutes after the reinforcement training. Before the testing, the electric grid shock floor was removed from the ‘Y’ maze, so the mice did not experience foot shock during the test session. The activity of the mice in the ‘Y’ maze was evaluated in blocks of three trials and each trial was 15 minutes. In the first trial, the mice were allowed to explore the arena without any light stimulation and assessed the baseline behavior. The time spent in each arm was calculated to evaluate arm preferences after the foot shock training session. For the following trials, the Light zone and Safe zone were selected, as indicated previously. The time spent in each arm was calculated by tracking the animal’s movement in the ‘Y’ maze using Any-maze video tracking software (Stoelting, IL, USA). A heatmap was also generated for each trial which displays the amount of time mice spent in different parts of the arena. A range of colors indicate the total time spent in the area with blue indicating the shortest and the red as the longest time.

### Statistical analysis

All statistical analyses were done by Graph Pad prism software (Graph-pad, San Diego, CA). Statistics are displayed as mean ± SEM. Paired t test was used for the comparison. Differences were determined significant when p< 0.05.

## Results

### Olfactory information changes with stimulus duration

Duration is an important input parameter of sensory stimuli [21] and the temporal properties of the stimulus are key for obtaining necessary information from the environment [10,11]. To test whether stimulus duration influence the olfactory identity, we optically stimulated ChR2 expressing OSNs in olfactory bulb and assessed the behavior in response to light stimulation of 10 or 25 ms duration. After training, we tested the Light zone avoidance response to OB stimulation, linked to the foot shock. First, we tested the mouse baseline behavior and calculated the time spent in each arm by allowing the mice to freely explore the arena. The aim of the baseline behavior analysis was to confirm if the mice have a preference to a particular arm of the ‘Y’ maze after the foot shock training. Our baseline data show that mice spent almost equal amounts of time in both arms of the ‘Y’ maze (Left zone - 469.3 ± 26.73 s, Right zone-430.7 ± 26.73 s, P = 0.50, Figure 2A, Movie S1). Then, we stimulated the OB with 25 ms duration light pulses with an interval of 150 ms. We found that, mice avoided the Light zone during 25 ms OB stimulation and spent most of their time in the Safe zone (Light zone - 94.75 ± 19.47 s, Safe zone - 805.3 ± 19.47 s, P = <0.0001, Figure 2B, 2C, Movie S2, n = 6). We also tested whether the avoidance response to the light stimulation was equally probable in both arms of the ‘Y’ maze. To confirm this, in some trials we delivered light stimulation when the mice reached the Safe zone. We found that, mice avoided the Safe zone during the light stimulation, indicating that the avoidance response is clearly linked to the olfactory information, but not to spatial information.

**Figure 2.**
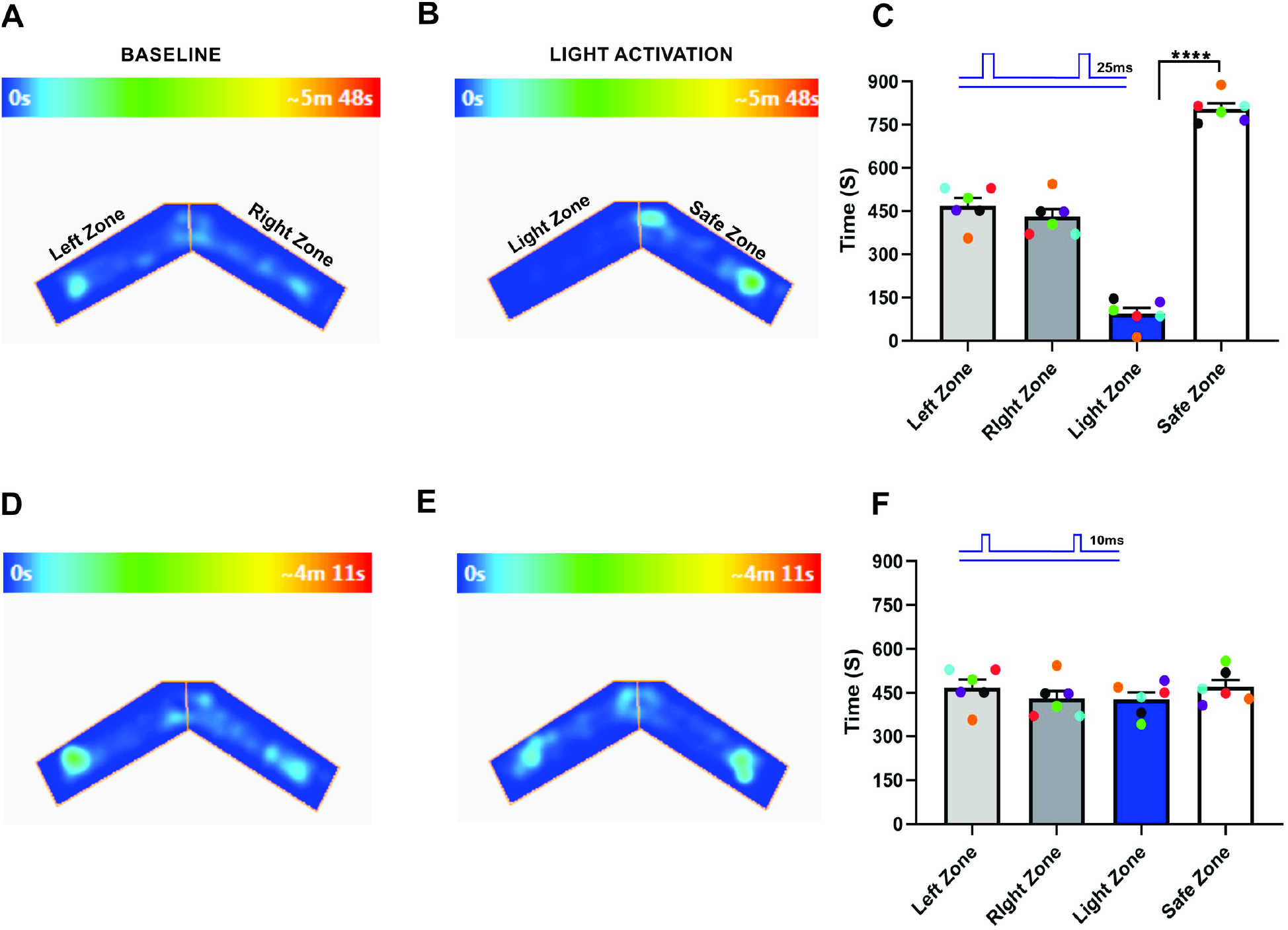
Stimulus duration changes the odor identity. A, B, an example of heat-map showing animal’s position in Y maze during baseline (A) and 25 ms light stimulation (B). C, Average amount of time explored in each zone in baseline and 25 ms light stimulation trials. D, E, Heat-map of mouse position during baseline (D) and 10 ms light stimulation (E). F, Average amount of time spent in each zone in baseline and 25 ms light stimulation trials.(****P<0.0001, n=6 animals, Heat map generated by ANY-maze version 5.2, https://www.anymaze.co.uk/index.htm).

Next, we stimulated the OB with 10 ms duration light pulses. Here, we found that the time spent in each arm during the baseline behavior trial (Left zone - 469.3 ± 26.73 s, Right zone - 430.7 ± 26.73 s, P = 0.50, Figure 2D) and the light activation trial (Light zone - 428.5 ± 23.26 s, Safe zone - 471.5 ± 23.26 s, P = 0.40, Figure 2E) were almost equal, indicating that the mice did not identify the foot-shock linked olfactory information. Thus, they did not avoid the Light zone during 10 ms OB stimulation (Figure 2F, Movie S3). In agreement with previous study in drosophila [22], our results suggest that duration of the stimulus changes the olfactory information.

### Bilateral input duration alters the unilateral olfactory identity

Odorant identity is one of the significant challenges for the olfactory system. Olfactory perception and behaviors are mainly depending on the ability to identify an odor across a wide variety of odor mixtures [23]. Previous studies report that bilateral olfactory input enhances the navigation and chemotaxis behavior[14,24]. To assess if the duration of the bilateral olfactory information influenced in olfactory identity, we stimulated each OB with different stimulus duration on unilaterally trained mice and observed if the mice can identify the foot-shock linked olfactory information from the synchronized bilateral OB stimulation. We found that during the baseline behavior trial, mice visited both arms equally (Left zone - 434.7 ± 58.81 s, Right zone - 465.3 ± 58.81 s, P = 0.80, Figure 3A), and when we synchronously stimulated each OB with 50 and 10 ms simultaneously, mice avoided the Light zone (Light zone - 91.08 ± 20.30 s, Safe zone - 808.9 ± 20.30 s, P = <0.0001, Figure 3B, 3C). Next, we synchronously stimulated each OB with 50 and 25 ms respectively, Here, we found that the time spent in each arm during the baseline behavior trial (Left zone - 450.3 ± 23.74 s, Right zone - 449.7 ± 23.74 s, P = 0.99, Figure 3D) and the light activation trial (Light zone - 487.4 ± 29.99 s, Safe zone - 412.6 ± 29.99 s, P = 0.27, Figure 3E, 3F) were almost equal, indicating that the mice did not identify the foot-shock linked olfactory information, thus, they did not avoid the Light zone. Our results suggest that, mice integrate the olfactory information from the contralateral OB during synchronized bilateral stimulation at longer stimulus duration and perceive it as a different olfactory information, but during the shorter stimulation, identity of the olfactory information remains same. Together, these results suggest that, duration of the bilateral input influences the olfactory identity.

**Figure 3.**
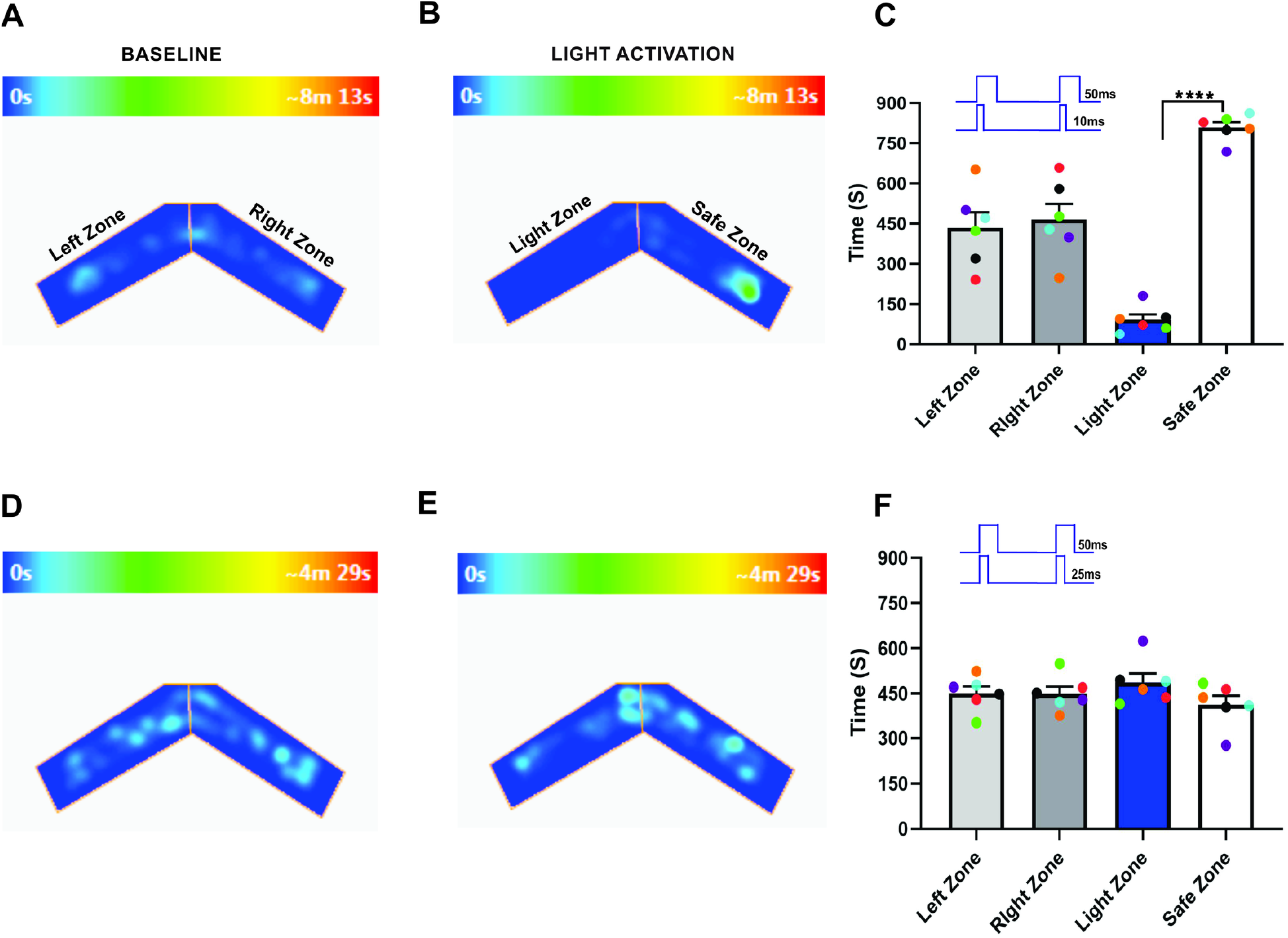
Bilateral input duration changes the unilateral olfactory identity. A, B, an example of heat-map showing animal’s position in Y maze during baseline (A) and dual OB (50 - 10 ms) stimulation (B). C, Average amount of time explored in each zone in baseline and dual OB (50 - 10 ms) stimulation trials. D, E, Heat-map of mouse position during baseline (D) and dual OB (50 - 25 ms) stimulation light stimulation (E). F, Average amount of time spent in each zone in baseline and dual OB (50 - 25 ms) stimulation trials. (****P<0.0001, n=6 animals).

To confirm this, we performed light stimulation and foot-shock avoidance training on a new group of animals (n=6). Here, we trained the mice with light stimulation to both olfactory bulbs simultaneously with 50 ms duration on each OB and paired it with foot shock. After the training, we stimulated each OB with different stimulus duration and tested the response of mice in the Light zone avoidance test. Our results show that when each olfactory bulbs were synchronously stimulated with 50 and 25 ms respectively, mice avoided the Light zone, indicating that mice can identify the bilaterally synchronized olfactory information linked to the foot-shock (Left zone - 455.8 ± 33.16 s, Right zone - 444.2 ± 33.16 s, P = 0.87, Light zone - 88.08 ± 19.28 s, Safe zone - 811.9 ± 19.28 s, P = <0.0001, Figure 4A, 4B, 4C, Movie S4). We then tested to determine if the mice can identify the foot-shock linked olfactory information when exposed to the stimulus duration of 50 and 10 ms respectively. We found that mice did not avoid the Light zone and continued to stay in the Light zone (Left zone - 468.5 ± 35.70 s, Right zone - 431.5 ± 35.70 s, P = 0.63, Light zone - 468.3 ± 26.24 s, Safe zone - 431.8 ± 26.24 s, P = 0.52, Figure 4D, 4E, 4F, Movie S5), suggesting that mice did not identify the bilaterally synchronized OB information. This confirms our previous results, that the bilateral input duration influences the olfactory identity.

**Figure 4.**
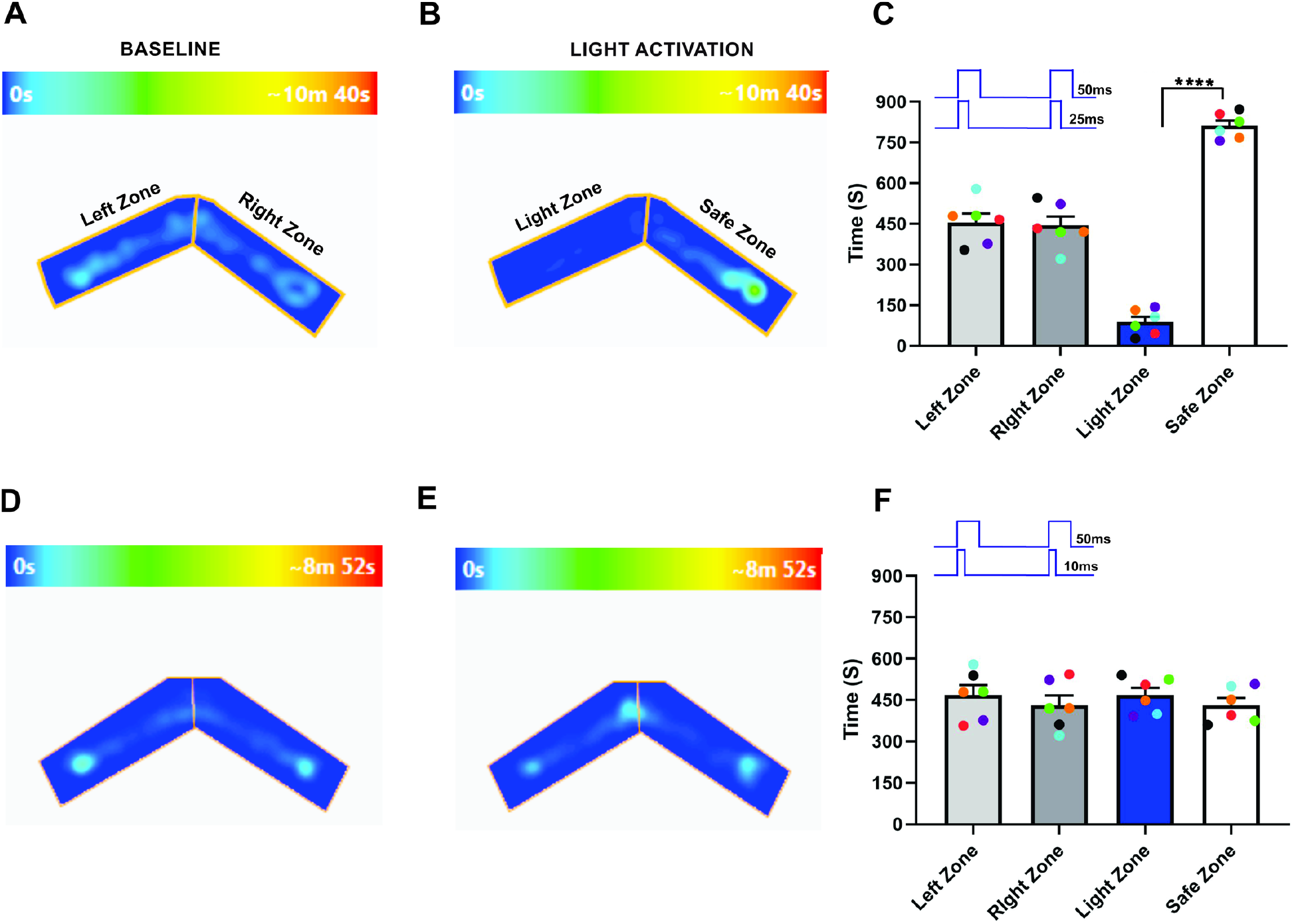
Stimulus duration influences the bilateral olfactory identity. A, B, an example of heat-map showing animal’s position in Y maze during baseline (A) and dual OB (50 - 25 ms) stimulation (B). C, Average amount of time explored in each zone in baseline and dual OB (50 - 25 ms) stimulation trials. D, E, Heat-map of mouse position during baseline (D) and dual OB (50 - 10 ms) stimulation (E). F, Average amount of time spent in each zone in baseline and dual OB (50 - 10 ms) stimulation trials. (****P<0.0001, n=6 animals).

Next, we checked if the olfactory information changes when both OB stimulated at same duration. Here, we synchronously stimulated both OB with 25 ms light pulses. Our results show that mice avoided the light zone during 25 ms light stimulation (Left zone - 430.7 ± 28.60 s, Right zone - 469.3 ± 28.60 s, P = 0.53, Light zone - 152.8 ± 34.03 s, Safe zone - 747.3 ± 34.03 s, P = 0.0003, Figure 5A, 5B, 5C, Movie S6). Followed by this, we simultaneously stimulated each OB with 10 ms light pulses and tested whether the mice detect the foot shock linked olfactory information during the short dual bulb stimulation. We found that, like baseline behavior, the mice spent almost equal amount of time in both arms of the ‘Y’ maze (Left zone - 474 ± 46.85 s, Right zone – 426 ± 46.85 s, P = 0.63, Light zone - 475.8 ± 45.38 s, Safe zone - 424.3 ± 45.38 s, P = 0.59, Figure 5D, 5E, 5F, Movie S7). Together, these results confirm that stimulus duration influences the olfactory identity.

**Figure 5.**
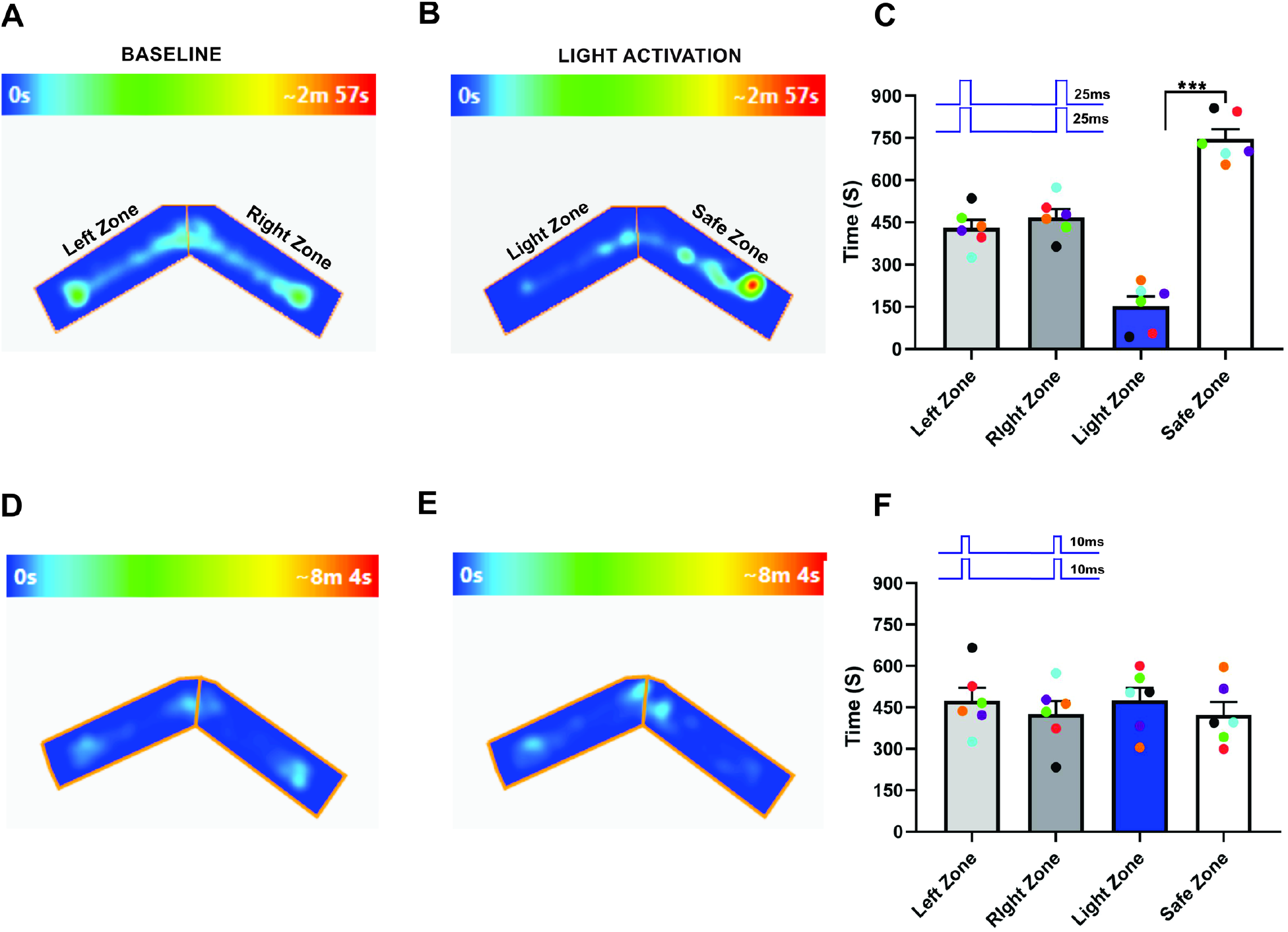
Bilateral olfactory identity. A, B, an example of heat-map showing animal’s position in Y maze during baseline (A) and dual OB (25 - 25 ms) stimulation (B). C, Average amount of time explored in each zone in baseline and dual OB (25 - 25 ms) stimulation trials. D, E, Heat-map of mouse position during baseline (D) and dual OB (10 - 10 ms) stimulation (E). F, Average amount of time spent in each zone in baseline and dual OB (10 - 10 ms) stimulation trials. (***P0.0003, n=6 animals).

Finally, to verify that the observed responses from the light stimulation were the result of activation of the ChR2 expressing neurons and not from the use of light as a visual cue, we used green light (540nm, output power, 20-22mw), which does not activate ChR2 and mice are relatively less sensitive to such long wavelength light [25–27]. During the green light stimulation, we did not observe significant behavioral difference from the baseline behavior (Left zone - 457.5 ± 51.09 s, Right zone - 442.5 ± 51.09 s, P = 0.89, Light zone - 463.8 ± 16.87 s, Safe zone - 436.2 ± 16.87 s, P = 0.45, Figure 6A-C), confirming that the mice did not use visual cues to perform the task. Together, our results demonstrate that animals might be detecting an olfactory information at shorter stimulus duration, but requires longer olfactory stimuli for identifying and generalizing an olfactory information.

**Figure 6.**
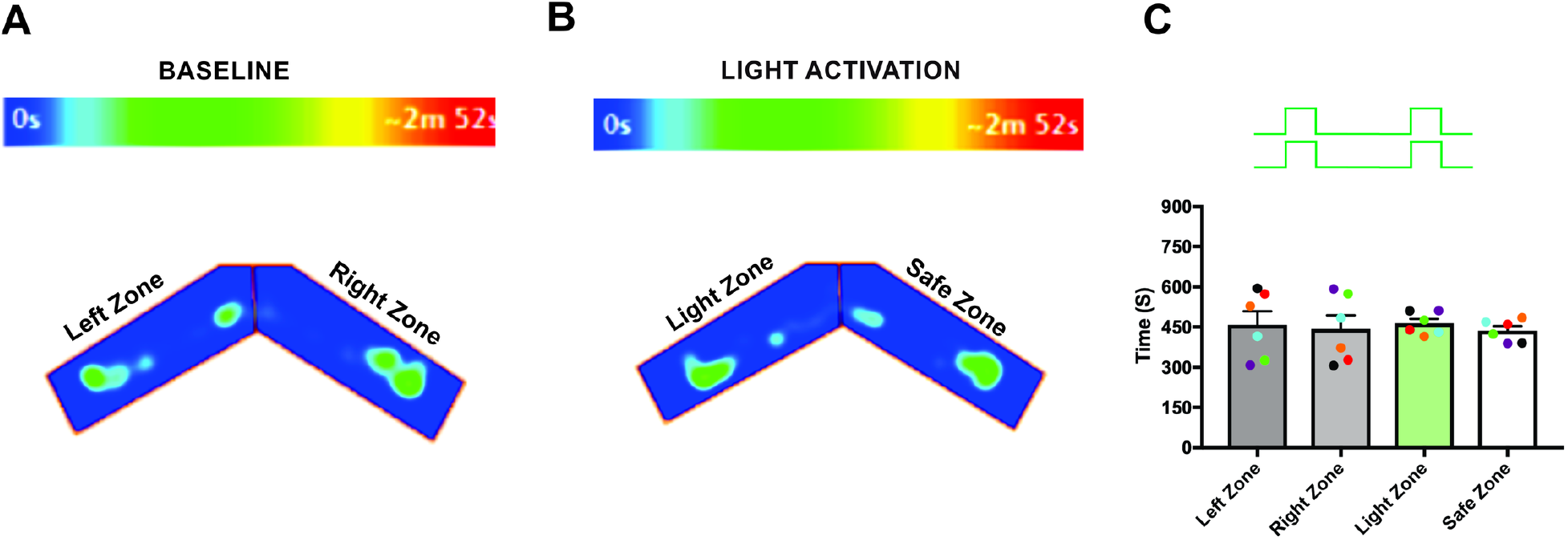
Green light stimulation did not activate the olfactory system of OMP-ChIEF mice A, B, Heat-map of mouse position during baseline (A) and green light stimulation (B). C, Average amount of time spent in each zone during green light stimulation.

## Discussion

Temporal properties of the stimuli are key for obtaining information from odor plumes in the environment[10,11]. In the olfactory system the role of stimulus timing is not well studied because of the difficulty of the precise control of odorant stimuli and the complex sniffing pattern of the animals. Previous studies have shown that such temporal coding could possibly transmit information regarding the direction and the concentration of an odor [5,15,21,28–31]. It is not well understood whether the temporal properties of an olfactory stimuli transmit any information about the identity of an olfactory information. Taking advantage of the transgenic mice, our study reveals that the duration of the olfactory stimuli carries information about the identity of an olfactory information. A previous study in drosophila have shown that flies can generalize an odor from a certain range of concentration, but when the concentration falls below certain threshold, the same odor will be detected as different [22]. It is possible that at lower concentrations, a smaller number of glomeruli are activated and when the concentration increases, a greater number of glomeruli get activated, which makes the odor generalization possible.

In rodents, identity and the generalization of an olfactory information can be influenced by the air flow rate, sniffing, complex interaction of the odorant molecule with the olfactory epithelium. Precise control of such parameters makes it hardly possible to study the role of stimulus duration in odor identity. Taking the advantage of optogenetics, Li et al., behaviorally showed that mice expressing ChR2 in olfactory sensory neurons can discriminate stimulus duration with a resolution of 10 ms [32]. Besides that, there are no reports available demonstrating the role of stimulus duration on olfactory identity. Previous studies in invertebrates has shown that they can encounter brief pulses of odor, and these pulses may contain valuable information about odor location, concentration, and odor identity [10,11,16]. In our study, we took the advantage of optogenetics to precisely control the activation of glomeruli expressing ChR2 to study the role of stimulus duration on olfactory identity. In accordance with studies in flies, our study provides evidence for the importance of stimulus duration on olfactory identity in vertebrates. Further research is required to fully understand how the stimulus duration influence the olfactory cortical neurons in encoding olfactory information for odor identity and odor localization. In natural environments, animals confront more complex problems such as, having to identify the quality, and complexity of an odor mixture. The use of suitable physiological and psychophysical paradigms will be a crucial step for further understanding the complexity of neural coding in the olfactory system.

## Notes

### Competing Interest Statement

The authors have declared no competing interest.

