## Supplemental Movie Legends for "The influence of stimulus duration on olfactory identity"

**Movie S1**. Mouse exploratory behavior

**Movie S2**. Unilateral foot shock trained mice avoid Light zone during 25 ms olfactory bulb Stimulation.

**Movie S3**. Unilateral foot shock trained mice did not avoid Light zone during 10 ms olfactory bulb stimulation.

**Movie S4**. Bilateral foot shock trained mice avoid Light zone during longer synchronized bilateral olfactory bulb stimulation (50 & 25 ms).

**Movie S5**. Bilateral foot shock trained mice did not avoid Light zone during shorter synchronous bilateral olfactory bulb stimulation (50 & 10 ms).

**Movie S6**. Bilateral foot shock trained mice avoid Light zone during synchronized bilateral olfactory bulb stimulation (25 & 25 ms).

**Movie S7**. Bilateral foot shock trained mice did not avoid Light zone during synchronized bilateral olfactory bulb stimulation (10 & 10 ms).
